# Tandem tension sensor reveals substrate rigidity-dependence of integrin molecular tensions in live cells

**DOI:** 10.1101/2020.01.24.918946

**Authors:** Anwesha Sarkar, Dana LeVine, Yuanchang Zhao, Keyvan Mollaeian, Juan Ren, Xuefeng Wang

**Author notes:** Correspondence and requests for materials should be addressed to X.W.

## Abstract

Response of integrin tensions to substrate rigidity is important in cell rigidity sensing but has not been confirmed. Current fluorescent tension sensors produce cellular force signals collectively resulted from integrin tension magnitude, tension dwell time, integrin density and activity, ligand density and accessibility, etc., making it challenging to monitor the absolute molecular force level of integrin tensions in live cells. Here we developed a tandem tension sensor (TTS) consisting of two coupled tension sensing units which are subject to the same tension and respond with different activation probabilities to the tension. Reported by fluorescence, the activation probability ratio of these two units solely responds to the force level of local integrin tensions, excluding the bias from other non-force factors. We verified the feasibility of TTS in detecting integrin tensions and applied it to study cells on elastic substrates. TTS unambiguously reported that integrin tensions in platelets decrease monotonically with the substrate rigidity, verifying the rigidity-dependence of integrin tensions in live cells.

## Introduction

Cellular forces at the cell-matrix interfaces are mainly transmitted by mechano-sensitive receptors, integrins [1–3]. These forces as biomechanical signals critically regulate a broad scope of cellular functions and physiological processes [4]. How cells produce and perceive these mechanical signals, termed mechanotransduction [5, 6], remains a central research interest in cell mechanobiology. Mechanotransduction is essential for cells to probe mechanical properties of local milieu. One remarkable trait of cells is that cells can generally sense the rigidity of their local environment and react accordingly. In the past, local rigidity has been shown to regulate a variety of cellular functions such as cell contraction, migration, proliferation and differentiation, and many physiological behaviors including cancer metastasis [7–9] and neuron development [10, 11]. However, how cells sense rigidity [12–15] is still not fully understood. Integrin tensions, the forces transmitted by integrin molecules, have been contemplated to participate in cell rigidity sensing. Intuitively, it is plausible that lower rigidity may relax cellular forces and reduce integrin tensions, and in turn impact the process of mechanotransduction. However, although bulk cellular forces have been demonstrated to be correlated with substrate rigidity [16, 17], integrin tensions as the minimal and fundamental mechanical signals in integrin signaling pathways, have not been monitored versus local rigidity. An important question remains unanswered: does rigidity alter the magnitude of integrin molecular tensions in adherent cells?

Despite the development of fluorescence-based molecular tension sensors [18–22] in recent years, it is still challenging to monitor the magnitude of integrin tensions. Targeting and binding to mechano-sensitive receptors, these surface-immobilized tension sensors undergo conformational change and convert local cellular force to fluorescence, enabling cellular force imaging at submicron resolution. In contrast to the significant improvement of resolution, these sensors do not exclusively respond to the magnitude of the integrin tensions. This is mainly because fluorescence signal induced by cellular force is not solely determined by molecular tension level, but contributed by many other factors: local integrin density, activity, ligand accessibility, force dwell time on individual integrins etc. Most of these factors are difficult to calibrate and control in cells. Moreover, molecular tension sensors can be activated by force in a large range of magnitude, making the resulted signal inaccurate in reporting the magnitude of integrin tensions or insensitive to the change of molecular tension level. Therefore, a tension sensor exclusively responding to the magnitude of integrin tensions will be valuable for the study of cell mechanotransduction and rigidity sensing.

In this work, we developed a tandem tension sensor (TTS) by coupling two force-reporting units in one construct. Each unit responds to a force by a probability function of the force level. Because these two units are coupled and subject to the same tension all the time, the ratio of their activation probabilities represented by the fluorescence intensities is therefore a function of the force level alone, excluding the impact of the local integrin density, activity and accessibility and other unforeseen factors. This ratiometric measurement eliminates the influence of all non-force magnitude factors and responds solely to the force level of integrin tensions. We constructed TTS and validated its performance using platelets, a type of anucleate mechano-sensitive cells which have robust adhesion force patterns [22]. We then incorporated TTS with elastic substrates and demonstrated that integrin tensions in both platelets and focal adhesions of adherent cells are positivity correlated with substrate rigidity in the range of 10kPa-1GPa.

## Results

### Structure of TTS

Previously, we developed integrative tension sensor based on double-stranded DNA (dsDNA) labeled with a dye-quencher pair [22, 23]. The mechanical separation of the dsDNA by a tension frees the dye from quenching, thus reporting the force with fluorescent signal. Here, a TTS construct consists of three DNA strands, with the upper strand conjugated with a blackhole quencher (BHQ2) and an integrin ligand RGD peptide, middle strand conjugated with an Iowa black red quencher (IAbRQ) and a Cy3, and lower strand conjugated with a biotin and Cy5. As shown in Fig. 1a, these three DNA strands are hybridized to form TTS. The biotin tag enables the surface immobilization of TTS through biotin-neutravidin interaction. The biotin is conjugated to the DNA with a triethyleneglycol spacer to reduce the steric hindrance during surface immobilization. Cy5 is quenched by IAbRQ and Cy3 is quenched by BHQ2, and RGD is presented to target integrins on cell membrane. The upper strand DNA forms an 18 base-paired (bp) dsDNA with a part of the middle-strand DNA, and the lower strand DNA forms a 21 bp dsDNA with another part of the middle-strand DNA. Five thymine nucleotides are positioned between the two dsDNAs as a spacer which rotates for about 180° so that the RGD is better presented to cells. When cells bind the RGD of the TTS and transmit the force to integrin-RGD bond, the integrin molecular tension may rupture either the 18 bp dsDNA or the 21 bp dsDNA of the TTS. The probability ratio of rupturing these two dsDNAs is evaluated by the Cy5 and Cy3 fluorescence signal intensities of the local TTS ensemble. In the extreme, at 0 force, the molecular ratio of activated Cy5-Cy3 would be the ratio between spontaneous dissociation rates of 21bp and 18bp dsDNAs. At a very high tension, the DNA unzipping rate becomes independent on the dsDNA length, and the Cy5-Cy3 molecular ratio is expected to be at 1 [24]. Next, we discuss the theoretical response curve of TTS to molecular tension in details.

**Fig. 1.**
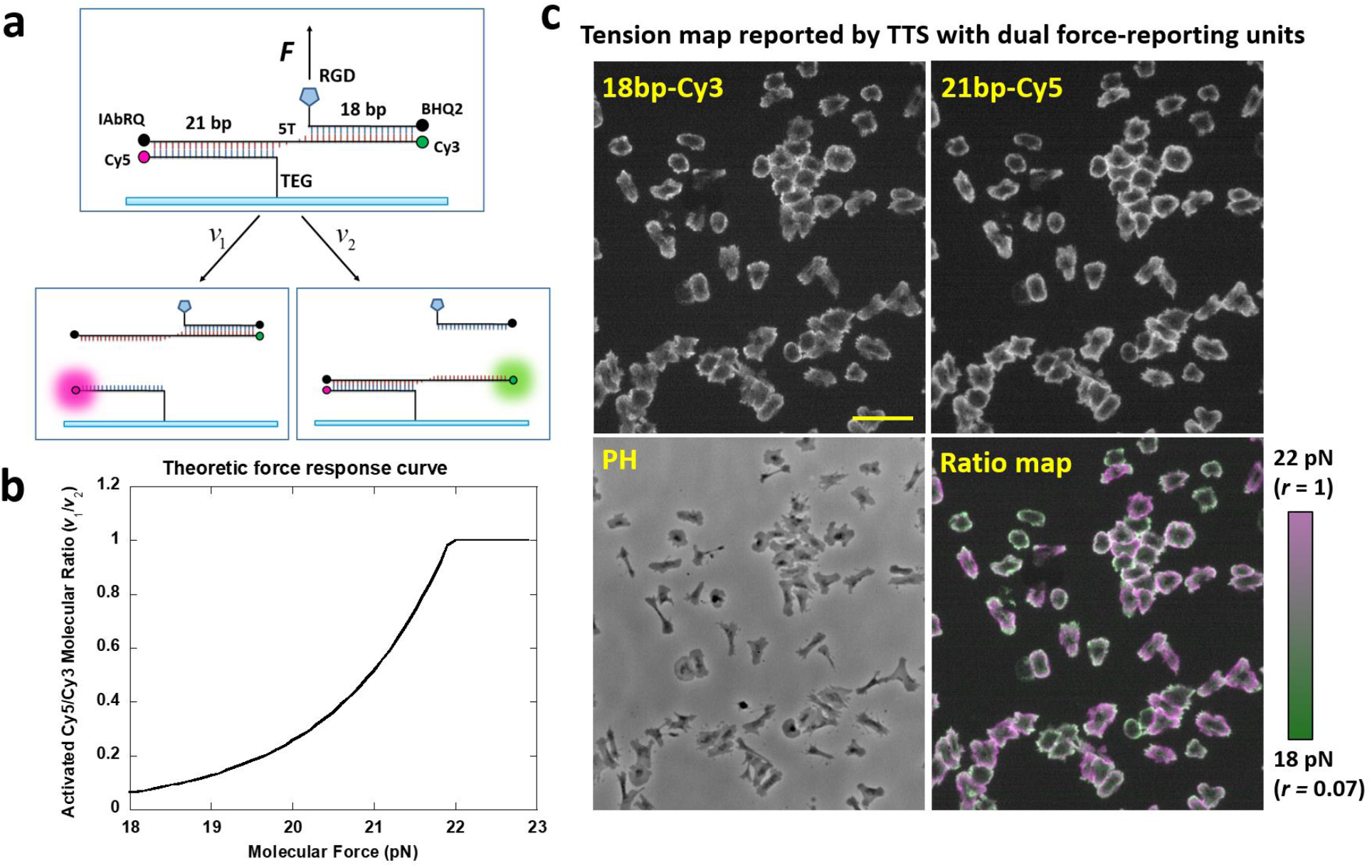
Tandem tension sensor (TTS) responds exclusively to integrin tension level by ratiometric measurement. **a**, The schematics of a TTS construct. TTS consists of a 21bp dsDNA and a 18bp dsDNA. Always subject to the same integrin tension, the two dsDNA dissociate at dissociation rates *v*_1_ or *v*_2_, respectively, leading to the activation of Cy5 or Cy3 with a probability ratio of *v*_1_/*v*_2_. **b**, Theoretical force response curve. Activation probability ratio of Cy5 and Cy3 is analytically derived and plotted against the molecular tension applied on the TTS. **c**, Test of TTS in reporting integrin tensions in platelets. Platelets were plated on a glass surface coated with TTS at a surface density 800/μm^2^. The local activations of Cy5 and Cy3 were imaged in two channels. Their activation probability ratio, i.e. the molecular ratio between the activated Cy5 and Cy3, was reported by the fluorescence intensities of Cy5 and Cy3 and visually appreciable by the color superposed images, with magenta indicating higher molecular tensions and green indicating lower ones. Under the imaging settings adopted in this work, the Cy5-Cy3 fluorescence intensity ratio is 1.5 times of activated Cy5-Cy3 molecular ratio.

### Theoretical response of TTS to molecular tension

Previous experiments and theoretical models of unzipping dsDNA by force have provided solid theoretical foundation for the force response of TTS. S. Cocco, et al. have derived the dissociation rate of dsDNA under a force. The dsDNA dissociation rate by an unzipping force is described as *v* = *v*_+_e^−*N*(2*g*(*f*)−*g*_0_)/*k_B_T*^ [24], where *v*_+_ is a constant independent of dsDNA length *N*, *k_B_* is the Boltzmann constant, *T* is temperature (*T* = 300*K* at room temperature) and g_0_ is the average free energy per base pair. *g*_0_ = −1.1*k_B_T* for a A-T pair and *g*_0_ = −3.5*k_B_T* for a C-G pair [24]. In TTS, the 21 bp dsDNA has 3 more C-G pairs than the 18 bp dsDNA. The free energy contributed by the unzipping force to each unzipped base is 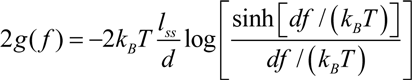 [24] and the coefficient factor 2 reflects the fact that two bases are unzipped for each base pair. In *g*(*f*), *l_ss_* = 0.56 nm is the contour length per base pair and *d* =1.5 nm is the Kuhn length of ssDNA [25, 26]. Based on this equation, we calculated the ratio of dissociation rates of 21 bp DNA and 18 bp DNA under the same unzipping force *f* as

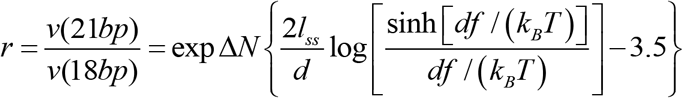

Note that *r* should be capped at 1, as the unzipping rate of a 21bp DNA is always less than that of a 18bp dsDNA. The theoretical force responsive curve of TTS is presented in Fig. 1b. This TTS construct is most sensitive to a force in the range of 18-22 pN which we found suitable for integrin tensions in adherent cells.

Here, we did not experimentally calibrate the force response of TTS because our emphasis is to monitor sensitivity of TTS to the alteration of integrin tensions in cells, rather than measuring the absolute value of the tension. The monotonic response of the Cy5-Cy3 ratio to force is a reliable conclusion derived from the DNA models backed up by substantial previous experiments and theory. TTS is expected to report the alteration of integrin tensions caused by external biochemical or mechanical cues with ultra-sensitivity and high specificity.

### TTS images molecular tensions in platelets

To verify the performance of TTS, we tested it in monitoring integrin tensions in platelets. Platelets are anucleate cells, mechanosensitive to local mechanics of the matrix [27]. With consistent morphology and adhesion force pattern [22], they are ideal model to test the cellular force response to substrate rigidity. Plated on the TTS immobilized on a glass surface, after 40 min, platelets produced force maps in both Cy3 and Cy5 channels (Fig. 1c). In either Cy3 or Cy5 channel, the fluorescence intensity of platelets is the collective result of integrin tensions and other non-force factors such as integrin density, integrin activation ratio and surface ligand accessibility. Therefore, the fluorescence intensity in either channel alone cannot reflect the magnitude of integrin tensions. However, the Cy5-Cy3 molecular ratio, representing the ratio of unzipping probabilities of 21bp and 18bp dsDNAs, is only determined by the integrin tension magnitude because the dual force-reporting units in TTS are consistently subject to the same force and non-force factors, therefore exclusively responding to integrin tension level. The Cy5-Cy3 molecular ratio is converted from Cy5-Cy3 fluorescence intensities (quantification process is shown in sFig. 1). Under the imaging settings in this work, Cy5-Cy3 molecular ratio is calculated by dividing Cy5-Cy3 fluorescence intensity ratio with a factor of 1.5 which is calibrated with a TTS construct without quenchers (sFig. 2). Cy5-Cy3 fluorescence intensity ratio can also be visually appreciated by the color in the merged images of Cy3 (green) and Cy5 (magenta) channels, with magenta indicating higher tension and green indicating lower tension. Therefore, Cy5-Cy3 fluorescence ratio serves as a marker indicating the average magnitude of local integrin tensions. The ratio map in Fig. 1c clearly shows variation of integrin tension level among platelets, as some ratio images of individual platelets exhibit magenta color and some exhibit greener color, showing that integrin tension levels are not uniform among these platelets. This is likely due to the different platelet adhesion time, as some platelets fell onto the surfaces earlier than others and exhibited higher force levels.

### TTS reveals that molecular tensions increase during platelet contraction

Platelets were previously shown to have two adhesion stages: initial spreading and later contraction. Platelet contraction is accompanied with stretched cell morphology and higher cellular force [28, 29]. Here, we tested if TTS can monitor the increase of cellular forces of platelets due to platelet contraction prompted by longer incubation time. In experiments, four aliquots from the same platelet solution were plated on four TTS surfaces (glass), respectively. The four samples were then fixed after incubation in an incubator for 10 min, 20 min, 40 min and 1 h, respectively. Because TTS is compatible with cell fixation, the ratio map is fixed and preserved at the desired time point. In Fig. 2a, four example ratio maps were presented. The cell surface density with earlier cell fixation time is lower because many platelets did not precipitate on the surface at the time. We quantified the average fluorescence intensity of individual platelets in both imaging channels. The Cy5 and Cy3 fluorescence intensities for individual platelets were quantified and plotted as a scatter plot in Fig. 2b. 100 platelets were analyzed for each incubation time. The force responses of platelets with different incubation times formed four distinct groups in the scatter plot. The Cy5-Cy3 molecular ratio is calculated for each platelet by dividing the Cy5-Cy3 intensity ratio with the pre-calibrated conversion factor 1.5. These molecular ratios were plotted in Fig. 2c. The Cy5-Cy3 molecular ratio reporting integrin tension level exhibits monotonic increase by platelet incubation time, suggesting that platelets with longer incubation time on a surface produce integrin tensions at a higher level. We further verified the correlation between platelet contraction and integrin tension level. Because platelets undergoing contraction have stretched morphology and larger cell shape index (CSI), we analyzed the CSI of platelets with 40 min incubation time and plotted Cy5-Cy3 molecular ratio against platelet CSI in Fig. 2d and sFig. 3. The result shows 0.68 correlation between CSI and Cy5-Cy3 molecular ratio, suggesting that TTS successfully reported a higher level of integrin tensions during platelet contraction.

**Fig. 2.**
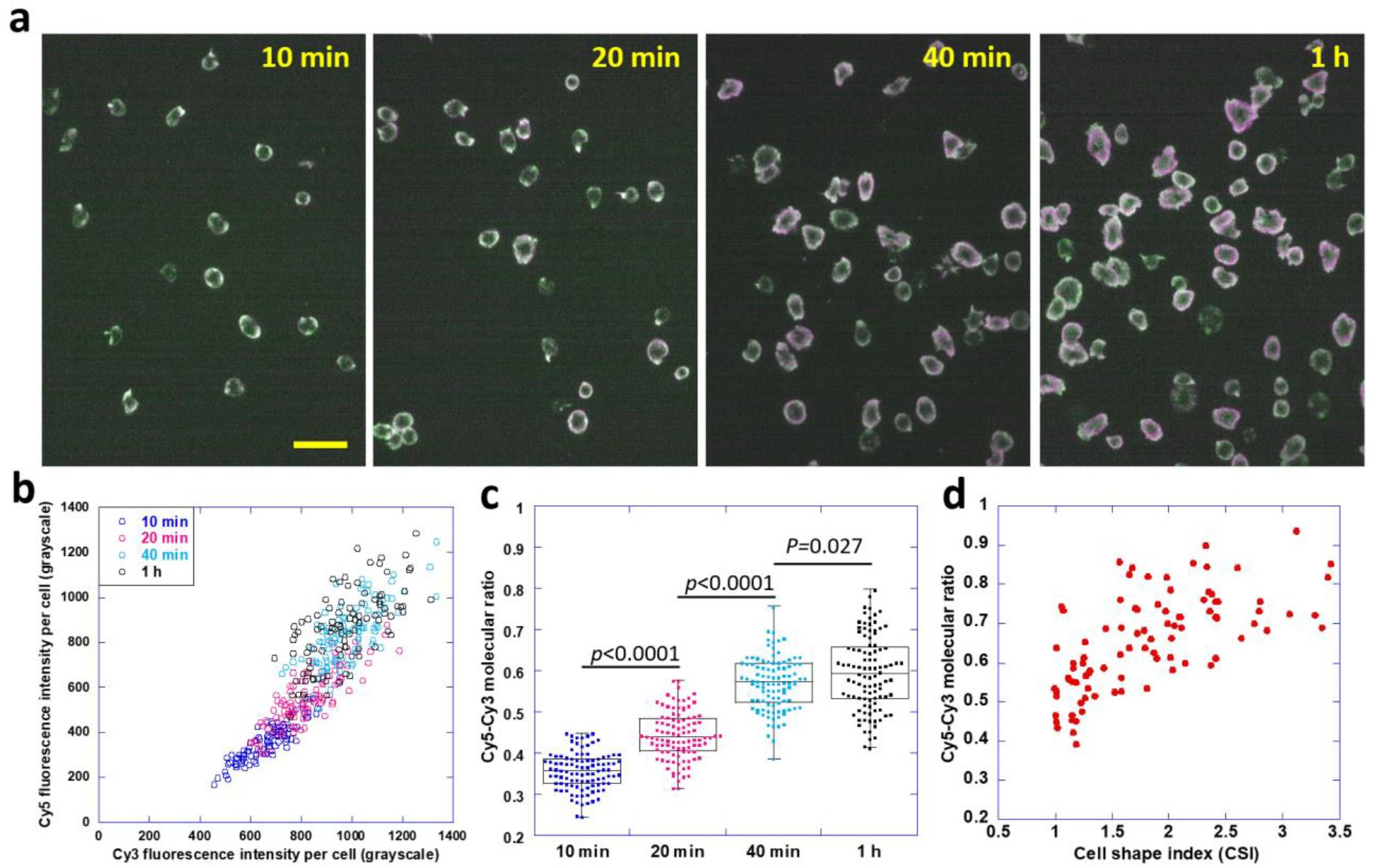
TTS reported integrin tensions varied by cell incubation time and cellular force inhibitors. **a**, TTS images of four aliquots of the same platelet sample which were incubated on TTS surfaces for 10 min, 20 min, 40 min and 1 h prior to imaging, respectively. Scale bar: 20 μm. **b**, a scatter plot of Cy5 and Cy3 fluorescence intensities (averaged by pixels) per platelet. 100 platelets were analyzed for each incubation time **c**, the molecular ratio of activated Cy5 and Cy3 per platelet. Molecular ratio is calculated by dividing Cy5-Cy3 fluorescence ratio by a pre-calibrated conversion factor, 1.5. **d**, Cy5-Cy3 ratio is correlated with cell shape index (CSI), with larger CSI indicating platelet contraction.

### TTS detects the decrease of molecular tensions due to myosin II inhibition

We verified that TTS reports higher molecular tensions during platelet contraction. It is known that platelet contraction requires actomyosin (Actin coupled with myosin II) which is a common source for integrin-transmitted cellular forces [30, 31]. Therefore, biochemically inhibiting actomyosin is expected to reduce integrin tensions in platelets. In order to test if TTS may report the change of intension tensions in platelet treated with actomyosin inhibitors, we performed TTS assays on platelets treated with three inhibitors, respectively: blebbistatin inhibiting myosin II, Y27632 inhbiting Rho-associated protein kinases (ROCK) and ML-7 inhibiting myosin light chain kinase (MLCK). ROCK is essential regulative protein in cell contraction and integrin tension generation in focal adhesions [32]. Inhibiting ROCK using Y-27632 abolishes integrin tension in focal adhesions [33]. MLCK is also involved in cell contraction by regulating myosin II function [34]. Previous studies show the both ML-7 and blebbistatin reduce bulk platelet forces but blebbistatin is more potent than ML-7. In Fig. 3a, the Cy5-Cy3 ratio maps are acquired and presented for the three samples and a control sample. All the three samples treated with inhibitors show greener color, indicating lower integrin tensions represented by lower Cy5-Cy3 ratios. The scatter plot of Cy5 and Cy3 intensities of individual platelets in Fig. 3b exhibits that four group of platelets are distinguishable from each other. Fig. 3c shows that Cy5-Cy3 molecular ratio became significantly lower in the three inhibited groups, suggesting that the Cy5-Cy3 ratios properly reflect the change of integrin tensions in platelets. Among the three inhibitors, blebbistatin exhibited the most potent effect in reducing the force level of integrin tensions (Fig. 3c).

**Fig. 3.**
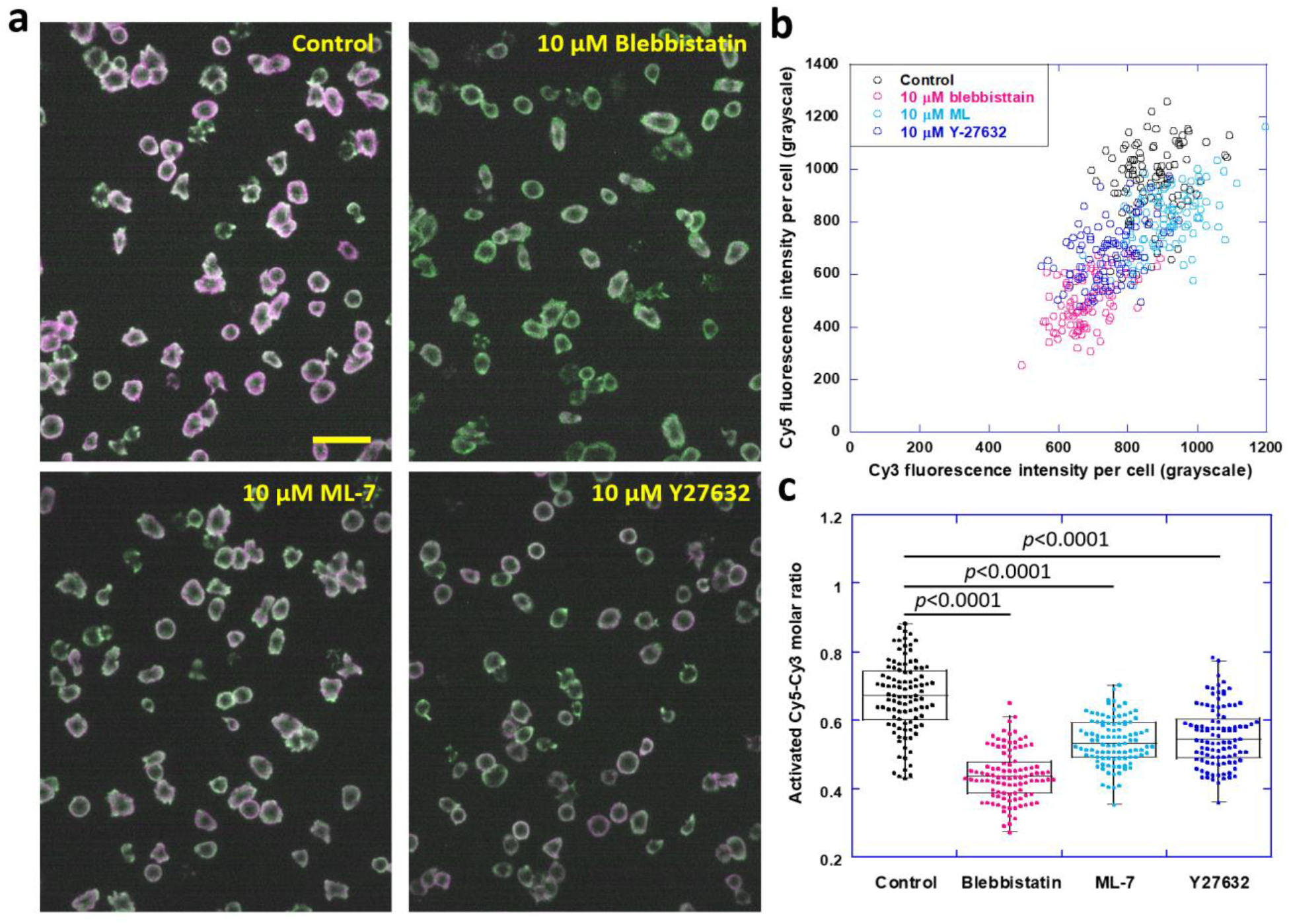
TTS reported integrin tensions altered by myosin inhibitor and ROCK inhibitor. **a**, TTS images of four aliquots of the same platelet sample which were treated with no inhibitor (control), blebbistatin, ML-7 and Y27632, respectively. Scale bar: 20 μm. **b**, a scatter plot of Cy5 and Cy3 fluorescence intensities per platelet. **c**, the molecular ratio of activated Cy5 and Cy3 per platelet.

### Integrin tensions in platelets monotonically decrease with substrate rigidity

With the experimental confirmation of TTS’s feasibility in reporting alteration of integrin tension level, we are ready to test whether the substrate rigidity dictates integrin tensions in live cells. Polydimethylsiloxane (PDMS) substrates of different rigidities were prepared by varying the volume ratio of base and crosslinker [35]. Substrates with Young’s modulus of 1.704 MPa, 213 kPa and 12 kPa were prepared and calibrated respectively (sFig. 4). TTS was coated on glass (1.7 GPa) and the three PDMS substrates with similar TTS surface densities which were calibrated in sFig. 5. Four portions of a same platelet solution were plated simultaneously on these substrates and imaged after 40 min incubation. It is qualitatively appreciable that the color of the ratio map changes from magenta to green when the substrate rigidity decreases, suggesting higher integrin tensions on more rigid surfaces (Fig. 4a). In Fig. 4b, the activated Cy5 and Cy3 intensities within 100 platelets were analyzed on each surface and plotted as a scatter plot. Platelets from four surfaces are clearly distributed in four separate groups in the plot. Cy3 and Cy5 intensities for single platelets are the strongest on glass surface and Cy5/Cy3 ratio represented by the slope is also the highest. The Cy5-Cy3 ratios of single platelets were calculated and presented in Fig. 4c which shows that force level of integrin tensions decreases monotonically with substrate rigidity, demonstrating a positive correlation between integrin tensions and substrate rigidity.

**Fig. 4.**
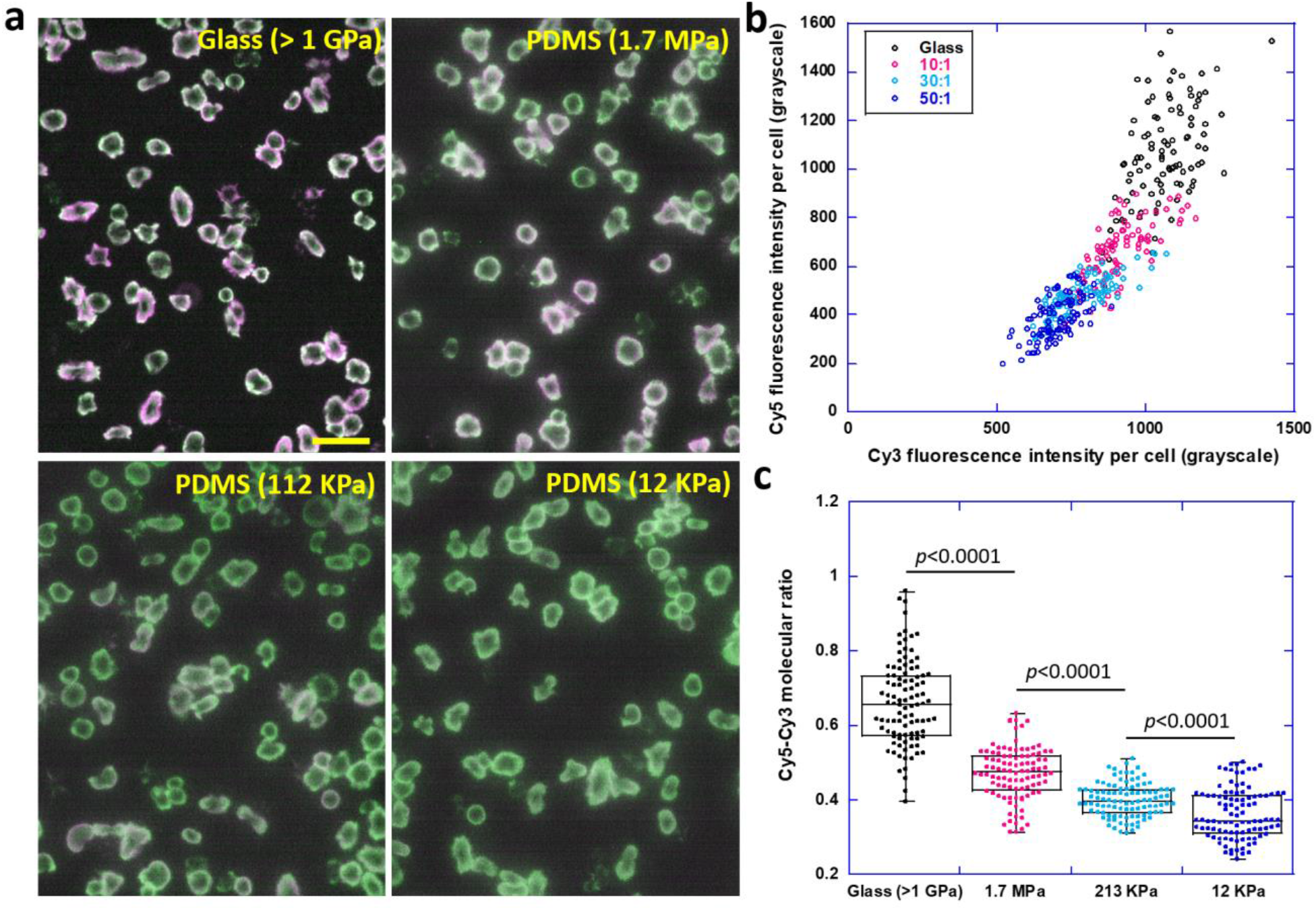
The force level of integrin tensions in platelets monotonically decrease by the rigidity of substrates. **a**, Ratiometric map of integrin tensions in platelets on TTS coated glass and PDMS. **b**, Scatter plot of Cy3 and Cy5 intensities of individual platelets. **c**, The molecular ratio of activated Cy5 and Cy3 per platelet versus substrate rigidity.

## Discussion

Numerous studies showed that matrix rigidity critically regulates cellular functions and impacts physiological processes such as stem cell differentiation [30, 36], cancer cell invasion [37]. Previously, platelets have been shown to exhibit the ability of rigidity sensing which is important for hemostasis [27]. However, how cells sense rigidity is not fully understood. The integrin molecular tension in cells is contemplated to affect the stability of molecular interactions and conformational change of mechanotransducive proteins, leading to the conversion of mechanical forces to biochemical signals [38]. Therefore, revealing the relationship between integrin tensions and substrate rigidity is important for the understanding of cell rigidity sensing.

To exclusively monitor the magnitude of integrin tensions, we developed TTS based on ratiometric force sensing. TTS consists of two coupled dsDNAs as dual force sensing units. The ratio of unzipping probabilities of these two dsDNAs reported by fluorescence is adopted as the indicator of the tension level exerted on the sensor. Previous studies of the kinetics of dsDNA unzipping by force provide a solid theoretical foundation for TTS performance. We also further demonstrated that integrin tensions reported by TTS did sensitively respond to platelet adhesion time and biochemical intervention, suggesting that TTS can reliably monitor the integrin tension level exerted by cells. Using TTS, we demonstrated that integrin tensions in platelets are dependent of substrate rigidity, therefore establishing the rigidity-dependence of integrin tensions in live cells.

TTS functions as a convenient approach to monitor force level of integrin tensions in cells at submicron resolution. It should be pointed out that TTS does not report integrin tension at the single molecule level but reflects the molecular tensions averaged at the local regions. However, TTS exclusively responds to the force level of integrin tensions, it can detect miniscule tension alteration caused by external or internal perturbation to cells. Overall, TTS is expected to probe the integrin tension levels with high resolution and ultra-sensitivity, providing a powerful approach for the study of cell mechanobiology at molecular tension level.

## Materials and Methods

### Tandem tension sensor (TTS) constructs

Tandem tension sensor was constructed by hybridizing three components: upper, middle and bottom strand single strand DNAs (ssDNA) as shown below in Table 1. All three ssDNAs were customized and purchased from Integrated DNA technologies, Inc. In order to synthesize the RGD (Arginylglycylaspartic acid, integrin ligand that can bind to both integrins αVβ3 and α5β1) linked upper strand, an amine tagged cyclic RGDfk was linked to BHQ2 tagged single-stranded DNA with thiol modification. Protocol of conjugating RGDfK to the DNA is included in this reference [39]. The middle strand was customized to be tagged with Iowa quencher (IAbRQ) and Cy3 fluorophore and the lower strand was conjugated to Cy5 fluorophore and biotin ensuring immobilization of TTS on a neutrAvidin coated surface using a biotin-neutrAvidin bond. These three strands were hybridized at a molar ratio of 1.4:1.2:1.0 (Top: Middle: Bottom) and annealed from 80°C to 25°C in PBS (phosphate buffered saline). The stocking concentration of TTS is 10 μM.

**Table 1.**
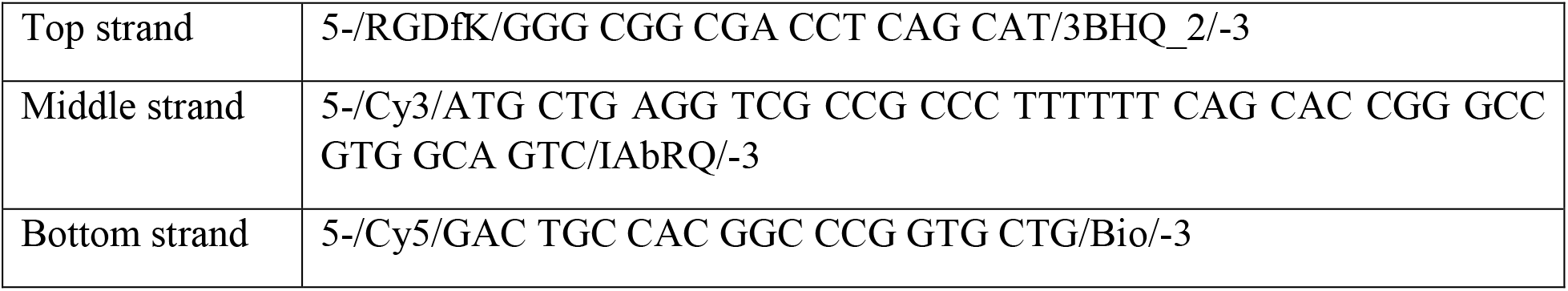
Three DNA strands for the construction of TTS.

### TTS immobilization on glass surfaces

TTS immobilization on glass starts with incubating a glass-bottom petridish (D35-14-1.5-N, Cellvis) with 200 μl solution with 100 μg/ml BSA-biotin (Biotinylated bovine serum albumin, 29130, Thermo Fischer scientific) and 5 μg/ml fibronectin in PBS at 4° C for 30 min. BSA-biotin subdues nonspecific adhesion of cells as well as serves biotin tags for the next layer of neutravidin. Fibronectin assists cell adhesion and minimize the impact of TTS rupture to cell functions. The sample was washed with 1×PBS (4°C) twice and next incubated with 100 μl solution of 100 μg/ml neutrAvidin (31000, Thermo Fischer scientific) in PBS at 4° C for 30 min followed by washing with PBS (4°C) twice. Next, the sample was incubated in 100 μl solution of 0.1 μM TTS in PBS at 4° C for 30 min followed by washing with PBS (4°C) twice. The sample was kept in a humidified chamber at 4° C.

The same coating protocol was applied to TTS coating on PDMS substrates.

### Preparation of PDMS substrates

In order to study integrin tensions in cells on PDMS substrates, polydimethylsiloxane (PDMS) samples were prepared at different volumetric ratio of base and crosslinker. PDMS is generally biocompatible and its mechanical properties are adjustable, making it an ideal platform for studying cellular behavior in response to substrate rigidity. Elastometer base and crosslinker (04019862, SYLGARD 184 silicone elastometer base and curing agent kit) were combined thoroughly with a spatula in a 1.5 ml microcentrifuge tube in ratios 10:1, 30:1 and 50:1, which result 1.7 MPa, 213 kPa and 12 kPa rigidities, respectively. 12 μl droplet of each mixture was dispensed on the glass well of 35 mm glass bottom dish with a 10 mm well (D35-10-1.5-N, Cellvis). The mixture was spread throughout the well with a pipette tip or spatula and the glass bottom dishes were kept on the bench top for 5 min for even coating of PDMS layer. Next, the samples were cured at 70°C for 24 h on a hot plate. This step was followed by shifting the dishes from hot plate to bench top and slowly cooling them down for an hour to bring the samples to room temperature.

### PDMS sample thickness calibration

Thickness calibration of the PDMS substrates was performed by brightfield imaging with a microscope with z axis quantification. Imaging was first focused on the top gel surface and then the bottom gel surface. The travel distance of the objective lens divided by the refractive index of PDMS (n=1.4) indicates the PDMS gel thickness. It was made sure that the gel thickness is at least 20 μm or above.

### PDMS stiffness calibration

Stiffness of the PDMS gel samples was calibrated using Atomic Force Microscopy (Bioscope catalyst AFM, Bruker corporation). The basic instrumentation of AFM comprises of a probe (a cantilever with sharp tip at the end), a cantilever deflection detecting system, a piezoelectric scanner, a sample holder and associated electronic control. Sample characterization was done by measuring the force of interaction between the cantilever tip and the sample surface. Peak Force QNM (quantitative nanomechanical mapping) mode was selected to perform elastic modulus measurement for all the PDMS samples. This mode is primarily built on the technologies of peak force tapping mode that can control the maximum applied forces on the substrate and thereby prevents damages to both cantilever and substrate. The set of parameters used for AFM measurement was peak force amplitude: 150 nm, peak force frequency: 2K Hz and scan rate: 0.497 Hz. PEAKFORCE-HiRs-F-A cantilever (Bruker AFM probes) with spring constant of 0.35 N/m was used for probing PDMS substrates. In our experiments, measurement of elastic modulus was performed by probing the cantilever tip on the polymerized gel samples and recording force vs distance curve. Considering the nature of the PDMS samples, the substrates will deform along with occurrence of elastic indentation according to Sneddon’s model of elasticity. Softer samples will also have more non-linear force curves compared to stiffer samples. This will result in lower cantilever deflection and flatter force curves. The retract curve of force vs tip-sample separating distance was fitted to the using the Sneddon model (this model was chosen keeping conical shape of the used cantilever tip in mind). Young’s Modulus of the substrates was directly derived from the software using the following Sneddon model equation:

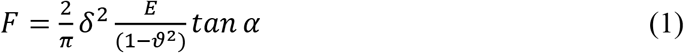

E in the above equation stands for Young’s modulus of the substrate, ϑ stands for Poisson’s ratio of the substrate in Equation (1). F represents the loading force on the substrate, δ stands for the indentation and α stands for opening angle of the conical cantilever tip. The peak force amplitude images of PDMS substrates of different stiffness and measured elastic modulus values were presented in Fig. S4.

### Extraction and preparation of Platelet-rich plasma (PRP)

Canine platelets were used for studying platelet force maps on TTS coated glass and PDMS substrates in this study. ISU Institutional Animal Care and Use Committee granted an approval for drawing the canine blood and the blood was drawn from healthy blood donor dogs or research dogs. Normally, a 3-mL syringe was used to collect 1.8 mL blood from a heathy dog’s cephalic vein. The syringe contained 0.2 ml Acid Citrate Dextrose or ACD buffer prior to blood collection in it. ACD buffer is a concoction of 41.6 mM citric acid, 136 mM glucose and 85.3 mM sodium citrate. The ACD buffer and blood combined solution was dispensed in a 15 ml falcon tube. The falcon tube already contained 2 mL buffered saline glucose citrate (BSGC). The BSGC buffer is a combination of 14 mM Sodium citrate, 11 mM glucose, 10 mM NaH_2_PO_4_, and 29 mM NaCl. This entire solution was centrifuged at 180g for 8 min without brake at room temperature (RT). The supernatant was collected in 1.5 ml eppendorf tubes as platelet-rich plasma (PRP). Post processing of PRP was always performed within 2 h of PRP preparation.

### Isolation of platelets for the Tandem tension sensor coated substrates

As activity of platelets is reduced with time, the prepared PRP must be used for TTS assay within 2 h after preparation. Platelets should also be purified from PRP before experiments as PRP possesses high level of anti-coagulant agents and fibrinogen. The purification process began by mixing 0.5 mL Buffer 1 (supplemented with 1 μM prostaglandin E1 and 0.02 U/ml apyrase) and 0.5 ml PRP in 1.5 ml eppendorf tube. Buffer 1 is a combination of 4.3 mM K_2_HPO_4_, 4.2 mM Na_2_HPO_4_, 113 mM NaCl, 24.3 mM NaH_2_PO_4_, and 5.5 mM Dextrose, pH 6.3. The mixture was centrifuged at 800 g for 8 min. The supernatant was pipetted out and the platelet pellet was gently re-suspended with serum-free Ham’s F-12 medium (HFL05-500ML, Caisson Laboratories).

### Activation and fixation of platelets

Platelets were re-suspended in serum free Ham’s F-12 medium with 100,000/μL concentration after purification and 10 μM Adenosine diphosphate (ADP) was added to the platelet solution for activation. Finally, platelets were plated on TTS coated substrates and incubated at for 40 min at 37°C for static imaging and for 6 min for real time imaging. For static imaging, all the samples were fixed using 4% paraformaldehyde (PFA) for 20 min at room temperature.

### HeLa cell culture, detaching and plating

HeLa cell lines were passaged every 72 hours on reaching 85% confluency. DMEM with L-Glutamine, 4.5g/L Glucose and Sodium Pyruvate (10-013-CVR, Thermo Fischer Scientific) supplemented 10% fetal bovine serum (FBS) and 1% streptomycin was used for culturing HeLa cells.

For cell plating, HeLa cells were first removed from bottom of culture flasks via a mild cell detaching solution in order to keep the cellular membrane proteins intact. The cell detaching solution contains 1× HBSS, 0.075% sodium bicarbonate, 10 mM HEPES (pH 7.6), 1.2 mM EDTA and H_2_O and the pH value of this solution was adjusted to 7.4 by adding NaOH or HCl. The existing medium in the culture flask was removed and cells were rinsed with 1 mL EDTA solution. Afterwards, 2 ml of EDTA solution was injected inside the culture flask. The flask was incubated for 10 min inside the incubator at 37°C. Afterwards, the cell solution was thoroughly pipetted multiple times and spun down for 3 min at 300 g. The supernatant was discarded and the cell pellet was re-suspended using 4 mL serum-free DMEM medium supplemented with 1% penicillin and streptomycin (PS). HeLa cells were plated on TTS coated substrates at a density of 10^6^/ml. HeLa cells were incubated for 1.5 hr at 37°C for static imaging.

### Cell force imaging on TTS surfaces

The imaging of platelets and HeLa cells were performed using a Nikon Ti-E fluorescence microscope. Phase contrast imaging was performed using 60× objective lens. Sola pad setting for both Cy3 and Cy5 Channels was kept at 30. Exposure time, used for Cy5 channel was 1 s; for Cy3 (TRITC) channel, it was 2 s. This imaging condition was maintained throughout our study for static imaging of fixed platelets and HeLa cells in this microscope.

### Immunostaining F-actin, and co-imaging of TTS, and F-actin

Following 1.5 h of incubation of HeLa cells at 37°C, fixation of the cells was performed using 4% paraformaldehyde (15710, Electron Microscopy Sciences) solution for 20 min at room temperature. This was followed by washing the cells 3 times with 1×PBS. Cell permeabilization was performed with 0.5% Triton X detergent at room temperature for 10 min. Afterwards, the cells were washed three times with 1×PBS. Next, 2.5 μg/ml dilutions of methanolic stocked Phalloidin-Alexa 488 solution (A22287, Invitrogen Molecular probes) was performed using 1×PBS. Duration of incubation in this solution was 20 min at room temperature. Finally, Cells were washed three times using 1×PBS. Imaging was performed immediately using 100× objective lens in Nikon Eclipse Ti-2 Total internal reflection fluorescence microscope (TIRFM).

## Acknowledgements

This work was supported by National Institute of General Medical Sciences (1R35GM128747) and National Science Foundation (1825724).

## Supplementary Figures

**sFig. 1.**
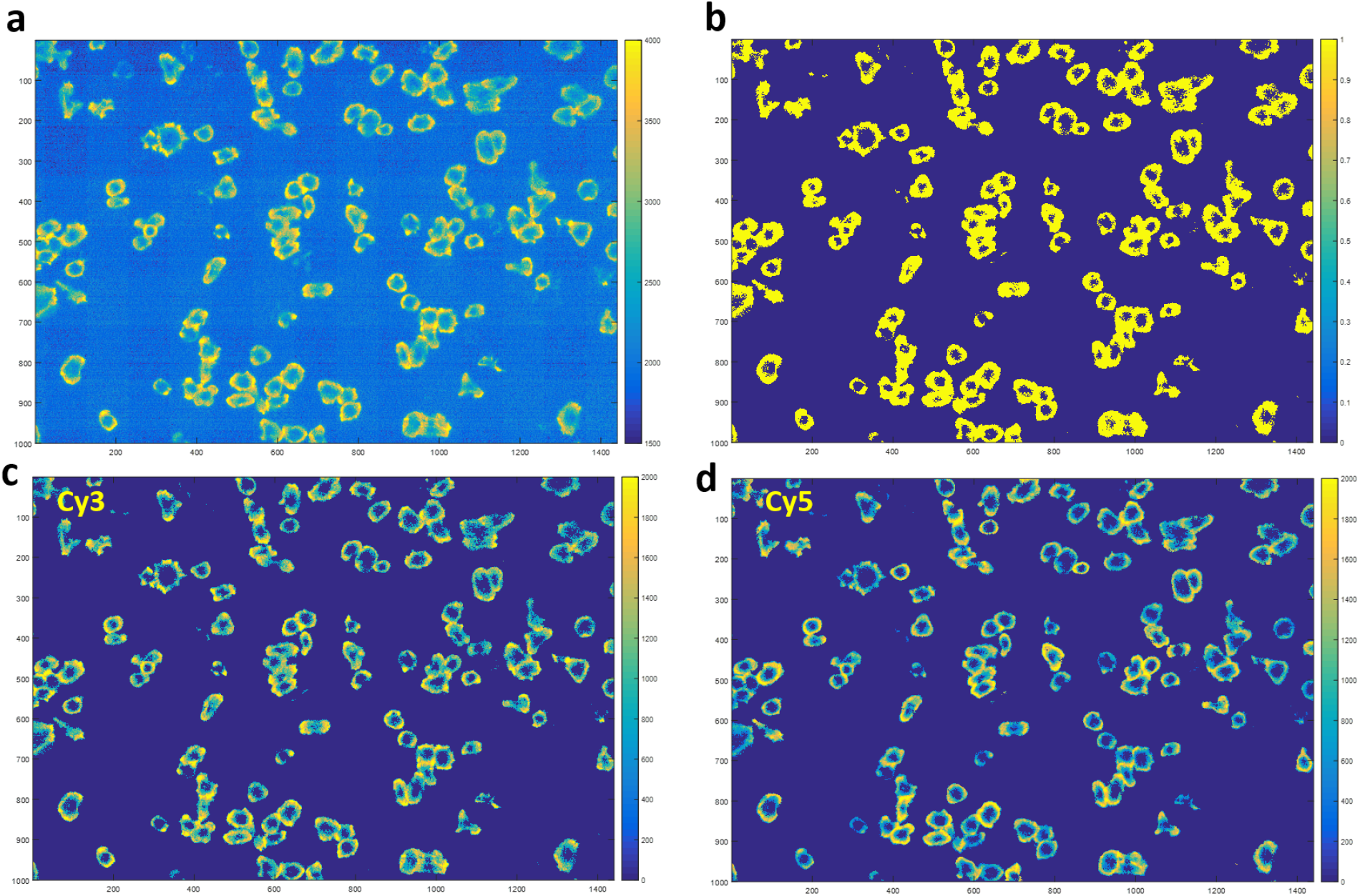
Calculation of Cy5-Cy3 fluorescence intensity ratio for individual platelets. **a**, Platelet force maps in Cy3 channel were analyzed to find the effective cell force regions, generating a mask to distinguish between force and background. **b**, Mask marking effective platelet force region (yellow region). **c-d**, Cy3 and Cy5 images multiplied by the mask and subtracted with the background grayscale value. Cy3 and Cy5 fluorescence intensities were calculated by summing the grayscale values of all pixels within individual platelets.

**sFig. 2.**
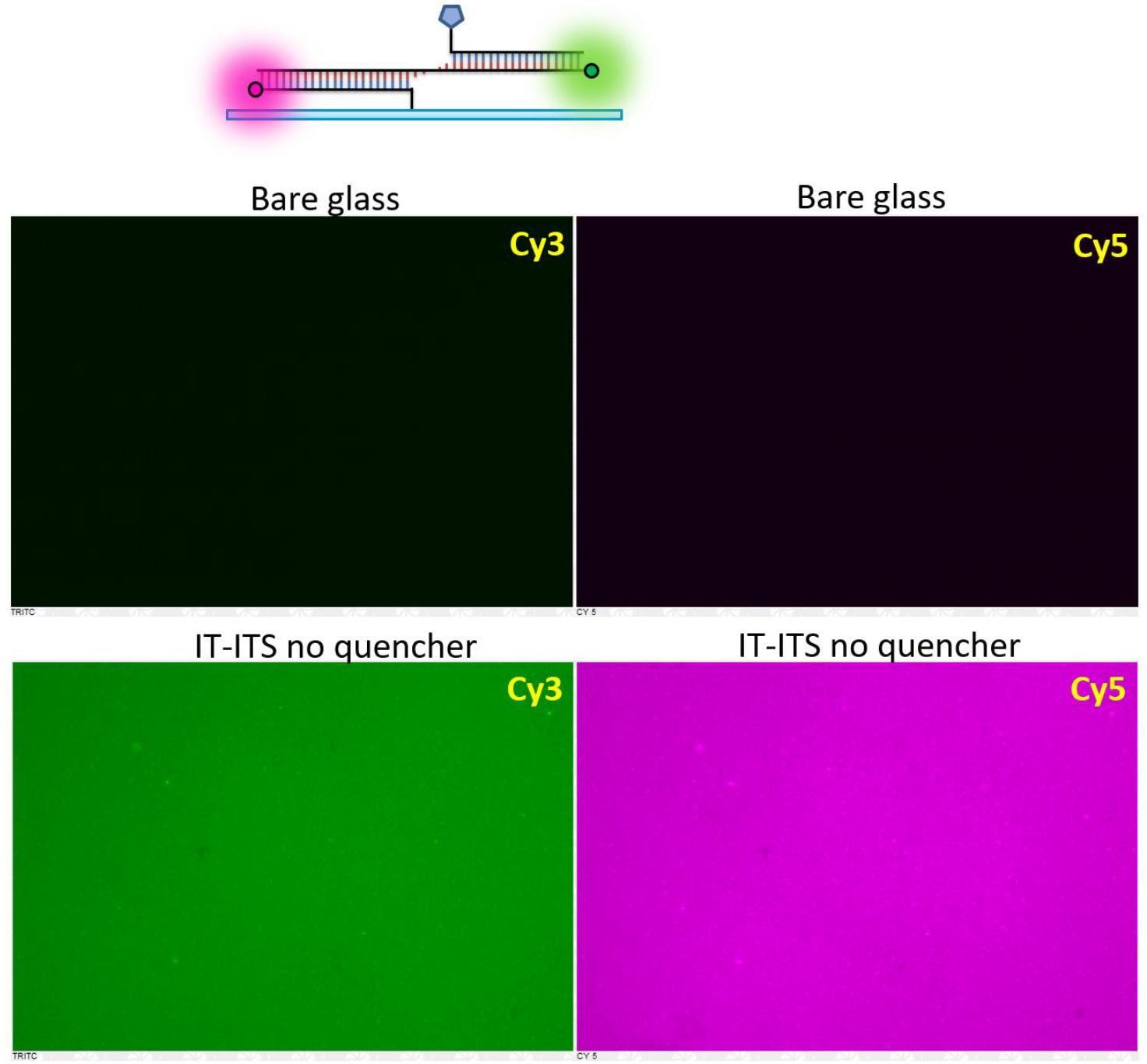
Calibration of fluorescence intensity ratio of Cy5 and Cy3. Cy5-Cy3 molecular ratio in this quencher-null TTS is 1:1. Such TTS produces Cy5-Cy3 fluorescence intensity ratio 1.5:1. Therefore, under current imaging settings, Cy5-Cy3 molecular ratio of regular TTS activated by cellular forces is Cy5-Cy3 fluorescence intensity ratio divided by a factor of 1.5.

**sFig. 3.**
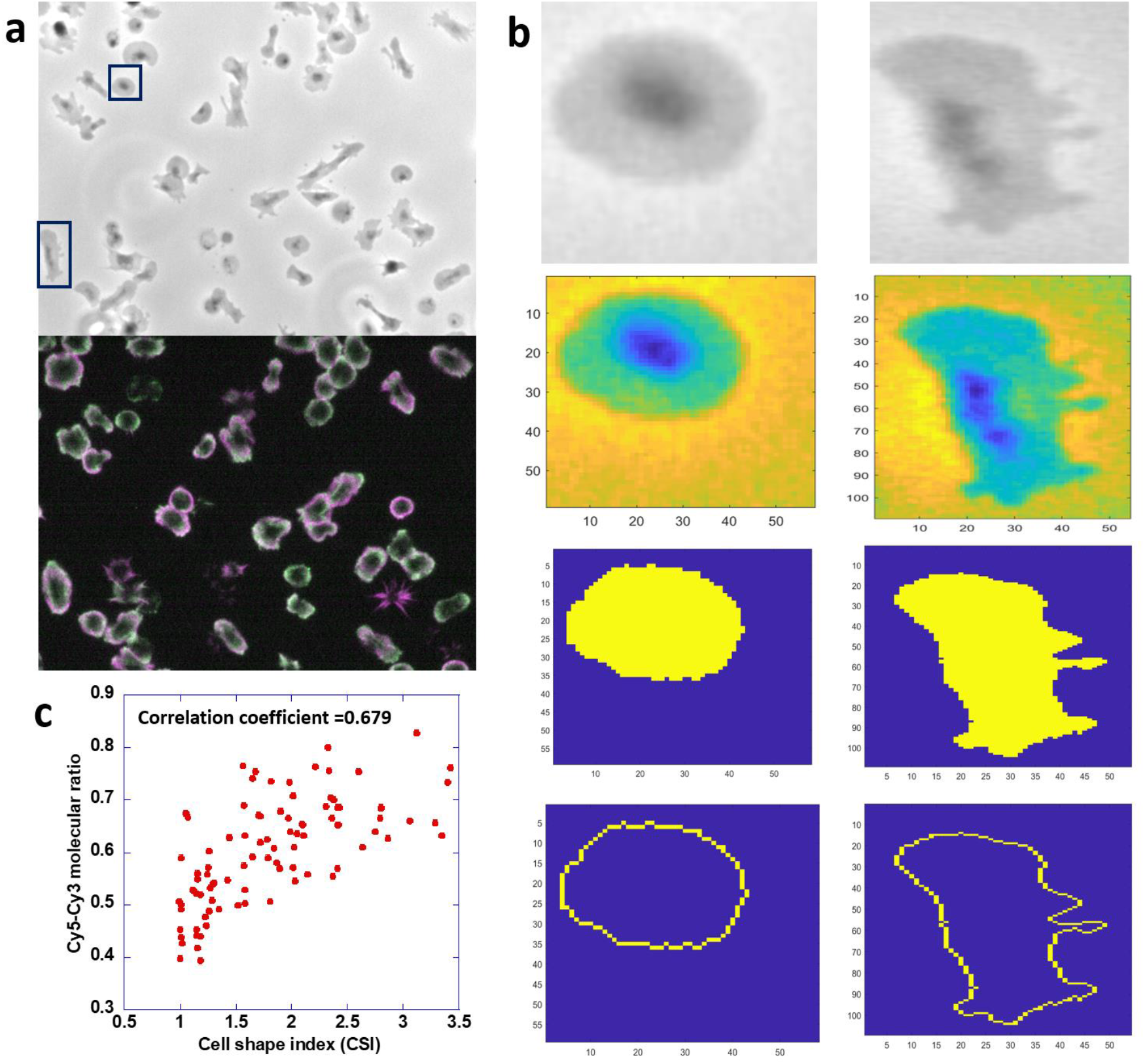
Correlation between Cy5-Cy3 molecular ratio and cell shape index of platelets. **a**, a phase-contrast image and a ratiometric force map for platelets. **b**, Matlab code was developed to find the platelet area (*A*) and contour length (*p*, perimeter) for individual platelets. Cell shape index is calculated as 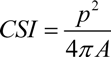. Minimum of CSI is 1, indicating a perfect circle. Larger CSI indicates more stretched cell morphology. **c**, Cy5-Cy3 molecular ratios of individual platelets were plotted against their CSIs.

**sFig. 4.**
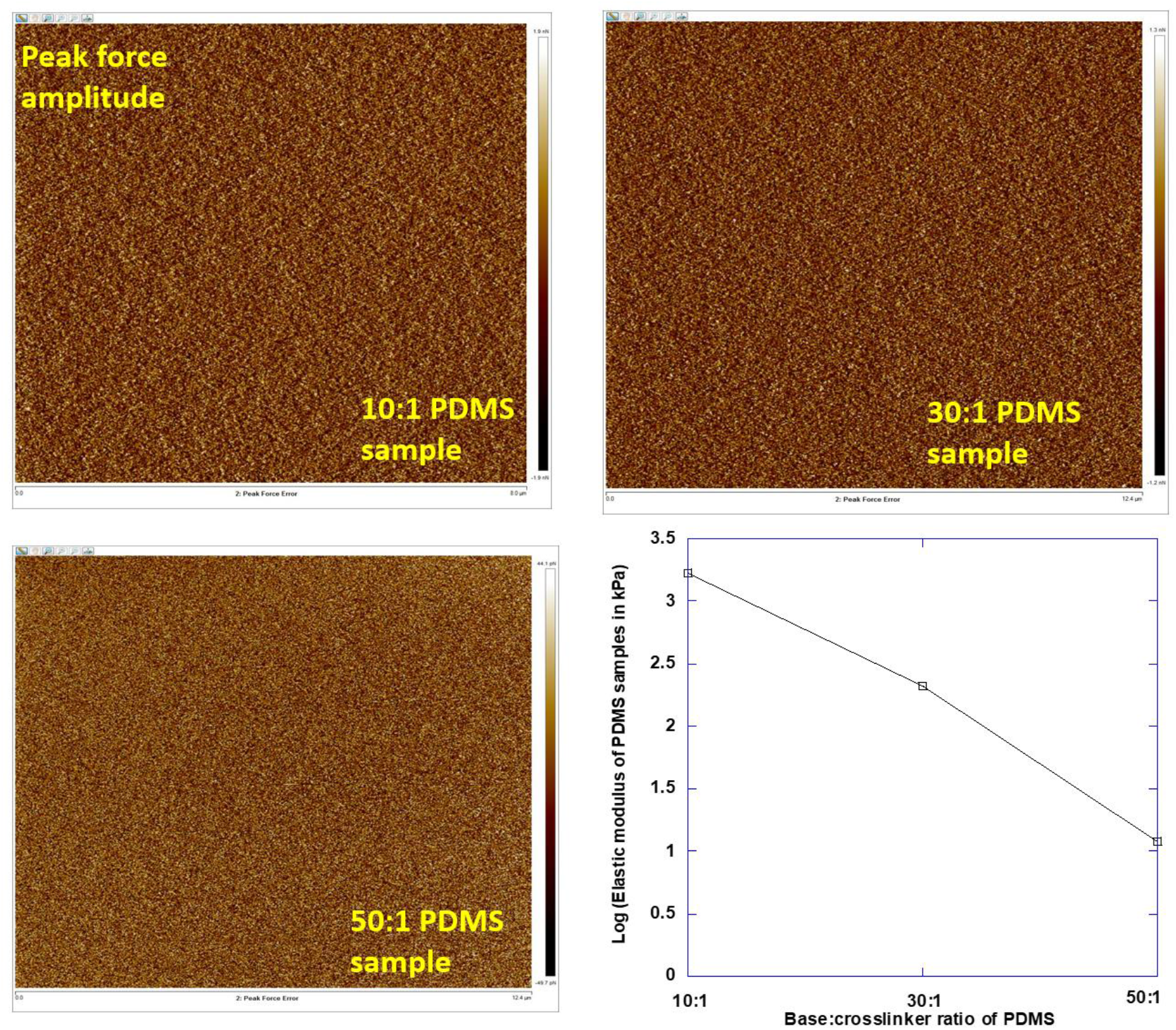
Calibration of elasticity of PDMS substrates. The AFM force maps were used to quantify the rigidities of PDMS substrates.

**sFig. 5.**
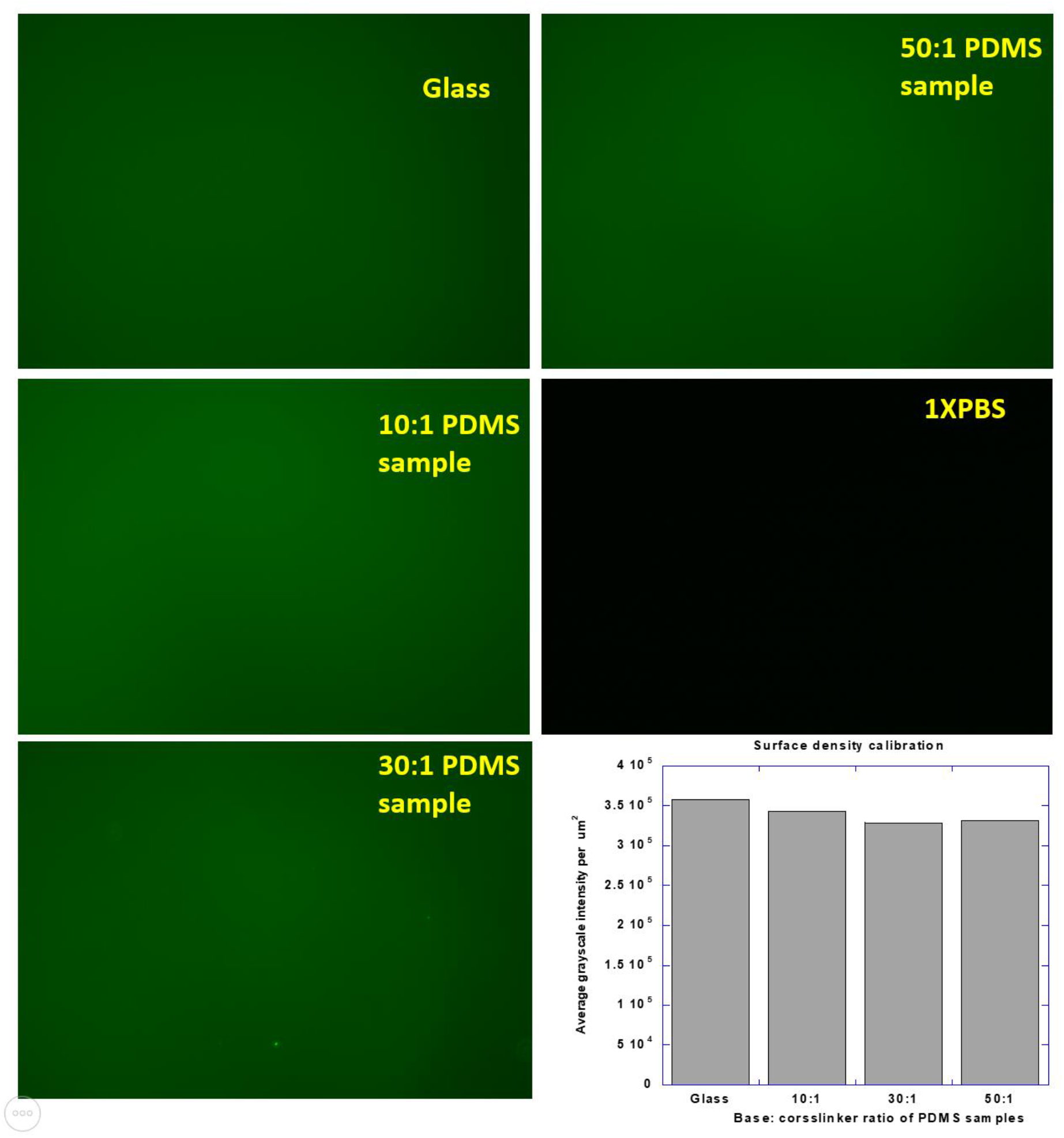
Calibration of relative TTS densities on glass and PDMS surfaces. Fluorescence images of quencher-null TTS on glass and three PDMS substrates were acquired, showing similar TTS coating densities on these substrates.

## References

1. Schwartz, M.A., Integrins and extracellular matrix in mechanotransduction. Cold Spring Harb Perspect Biol, 2010. 2(12): p. a005066.

2. Roca-Cusachs, P., et al., Clustering of alpha(5)beta(1) integrins determines adhesion strength whereas alpha(v)beta(3) and talin enable mechanotransduction. Proc Natl Acad Sci U S A, 2009. 106(38): p. 16245–50.

3. Katsumi, A., et al., Integrins in mechanotransduction. J Biol Chem, 2004. 279(13): p. 12001–4.

4. Vogel, V. and M. Sheetz, Local force and geometry sensing regulate cell functions. Nature Reviews Molecular Cell Biology, 2006. 7(4): p. 265–275.

5. Sun, Z., S.S. Guo, and R. Fassler, Integrin-mediated mechanotransduction. J Cell Biol, 2016. 215(4): p. 445–456.

6. Humphrey, J.D., E.R. Dufresne, and M.A. Schwartz, Mechanotransduction and extracellular matrix homeostasis. Nature Reviews Molecular Cell Biology, 2014. 15(12): p. 802–812.

7. Tilghman, R.W., et al., Matrix rigidity regulates cancer cell growth and cellular phenotype. PLoS One, 2010. 5(9): p. e12905.

8. Kostic, A., C.D. Lynch, and M.P. Sheetz, Differential matrix rigidity response in breast cancer cell lines correlates with the tissue tropism. PLoS One, 2009. 4(7): p. e6361.

9. Chin, L., et al., Mechanotransduction in cancer. Curr Opin Chem Eng, 2016. 11: p. 77–84.

10. Koch, D., et al., Strength in the periphery: growth cone biomechanics and substrate rigidity response in peripheral and central nervous system neurons. Biophys J, 2012. 102(3): p. 452–60.

11. Kostic, A., J. Sap, and M.P. Sheetz, RPTPalpha is required for rigidity-dependent inhibition of extension and differentiation of hippocampal neurons. J Cell Sci, 2007. 120(Pt 21): p. 3895–904.

12. Gupta, M., et al., Single cell rigidity sensing: A complex relationship between focal adhesion dynamics and large-scale actin cytoskeleton remodeling. Cell Adh Migr, 2016. 10(5): p. 554–567.

13. Moore, S.W., P. Roca-Cusachs, and M.P. Sheetz, Stretchy proteins on stretchy substrates: the important elements of integrin-mediated rigidity sensing. Dev Cell, 2010. 19(2): p. 194–206.

14. Jiang, G., et al., Rigidity sensing at the leading edge through alphavbeta3 integrins and RPTPalpha. Biophys J, 2006. 90(5): p. 1804–9.

15. Yip, A.K., et al., Cellular response to substrate rigidity is governed by either stress or strain. Biophys J, 2013. 104(1): p. 19–29.

16. Ghibaudo, M., et al., Traction forces and rigidity sensing regulate cell functions. Soft Matter, 2008. 4(9): p. 1836–1843.

17. Elosegui-Artola, A., et al., Mechanical regulation of a molecular clutch defines force transmission and transduction in response to matrix rigidity. Nature Cell Biology, 2016. 18(5): p. 540-+.

18. Morimatsu, M., et al., Molecular Tension Sensors Report Forces Generated by Single Integrin Molecules in Living Cells. Nano Letters, 2013. 13(9): p. 3985–3989.

19. Wang, X. and T. Ha, Defining single molecular forces required to activate integrin and notch signaling. Science, 2013. 340(6135): p. 991–4.

20. Zhang, Y., et al., DNA-based digital tension probes reveal integrin forces during early cell adhesion. Nat Commun, 2014. 5: p. 5167.

21. Blakely, B.L., et al., A DNA-based molecular probe for optically reporting cellular traction forces. Nat Methods, 2014. 11(12): p. 1229–32.

22. Wang, Y.L., et al., Force-activatable biosensor enables single platelet force mapping directly by fluorescence imaging. Biosensors & Bioelectronics, 2018. 100: p. 192–200.

23. Wang, X.F. and T. Ha, Defining Single Molecular Forces Required to Activate Integrin and Notch Signaling. Science, 2013. 340(6135): p. 991–994.

24. Cocco, S., R. Monasson, and J.F. Marko, Force and kinetic barriers to unzipping of the DNA double helix. Proc Natl Acad Sci U S A, 2001. 98(15): p. 8608–13.

25. Rief, M., H. Clausen-Schaumann, and H.E. Gaub, Sequence-dependent mechanics of single DNA molecules. Nature Structural Biology, 1999. 6(4): p. 346–349.

26. Smith, S.B., Y. Cui, and C. Bustamante, Overstretching B-DNA: the elastic response of individual double-stranded and single-stranded DNA molecules. Science, 1996. 271(5250): p. 795–9.

27. Qiu, Y.Z., et al., Platelet mechanosensing of substrate stiffness during clot formation mediates adhesion, spreading, and activation. Proceedings of the National Academy of Sciences of the United States of America, 2014. 111(40): p. 14430–14435.

28. Lam, W.A., et al., Mechanics and contraction dynamics of single platelets and implications for clot stiffening. Nature Materials, 2011. 10(1): p. 61–66.

29. Myers, D.R., et al., Single-platelet nanomechanics measured by high-throughput cytometry. Nature Materials, 2017. 16(2): p. 230–235.

30. Samuel, M.S., et al., Actomyosin-Mediated Cellular Tension Drives Increased Tissue Stiffness and beta-Catenin Activation to Induce Epidermal Hyperplasia and Tumor Growth. Cancer Cell, 2011. 19(6): p. 776–791.

31. Wozniak, M.A. and C.S. Chen, Mechanotransduction in development: a growing role for contractility. Nature Reviews Molecular Cell Biology, 2009. 10(1): p. 34–43.

32. Provenzano, P.P. and P.J. Keely, Mechanical signaling through the cytoskeleton regulates cell proliferation by coordinated focal adhesion and Rho GTPase signaling. Journal of Cell Science, 2011. 124(8): p. 1195–1205.

33. Wang, X.F., et al., Integrin Molecular Tension within Motile Focal Adhesions. Biophysical Journal, 2015. 109(11): p. 2259–2267.

34. Goeckeler, Z.M. and R.B. Wysolmerski, Myosin Light-Chain Kinase-Regulated Endothelial-Cell Contraction - the Relationship between Isometric Tension, Actin Polymerization, and Myosin Phosphorylation. Journal of Cell Biology, 1995. 130(3): p. 613–627.

35. Trappmann, B., et al., Extracellular-matrix tethering regulates stem-cell fate (vol 11, pg 642, 2012). Nature Materials, 2012. 11(8): p. 742–742.

36. Engler, A.J., et al., Matrix elasticity directs stem cell lineage specification. Cell, 2006. 126(4): p. 677–689.

37. Ulrich, T.A., E.M.D. Pardo, and S. Kumar, The Mechanical Rigidity of the Extracellular Matrix Regulates the Structure, Motility, and Proliferation of Glioma Cells. Cancer Research, 2009. 69(10): p. 4167–4174.

38. Schoen, I., B.L. Pruitt, and V. Vogel, The Yin-Yang of Rigidity Sensing: How Forces and Mechanical Properties Regulate the Cellular Response to Materials. Annual Review of Materials Research, Vol 43, 2013. 43: p. 589–618.

39. Zhao, Y.C., N.M. Wetter, and X.F. Wang, Imaging Integrin Tension and Cellular Force at Submicron Resolution with an Integrative Tension Sensor. Jove-Journal of Visualized Experiments, 2019(146).

